# No evidence for HPAI H5N1 2.3.4.4b incursion into Australia in 2022

**DOI:** 10.1101/2023.02.06.527378

**Authors:** Michelle Wille, Marcel Klaassen

## Abstract

There is an ongoing and profound burden of lineage 2.3.4.4b high pathogenicity avian influenza (HPAI) H5 on wild birds and poultry, globally. Herein we report the continued absence of HPAI in Australia from September – December 2022, in inbound migratory birds. Given the ever changing phenotype of this virus, worldwide studies on the occurence, or here absence of the virus, are of critical importance to understand the virus’ dispersal and incursion risk and development of response strategies.

## Main Text

The current high pathogenicity avian influenza (HPAI) H5 panzootic is having a profound impact on the poultry industry and wildlife (1). While lineage 2.3.4.4b is of current concern, HPAI H5 emerged in poultry in 1996 and has caused outbreaks in wild bird populations episodically since 2005 (2). The epidemiology of this virus has changed substantially with the emergence of new lineages, as exampled by Clade 2 viruses which caused the first wild bird mass mortality event at Qinghai Lake, China in 2005 (3). A novel lineage emerged in 2014 (2.3.4.4), which has diversified and caused substantial mortality, including mass mortality events, of wild birds in 2014, 2016, 2020-present, along with ongoing outbreaks in poultry in Eurasia and North America (2).

Understanding viral incursion risk following the emergence of novel lineages of HPAI with their own specific phenotype is of crucial importance in preventing incursion events, improving biosecurity to protect poultry, and responding to wild bird outbreaks. The viral incursion into North America in December 2021 was not detected until outbreaks occurred in poultry (4). Also the recent incursion into South America, in Nov 2022, was only detected following mass mortality events (5). Wild migratory waterfowl have been predominantly implicated in the re-occuring incursions into Europe and Africa (6). However, there are few migratory waterfowl linking the Nearctic and Palearctic, as well as North and South America, suggesting that the long-distance dispersal of lineage 2.3.4.4b HPAI may rely on additional bird groups but waterfowl [*e.g*. (7)].

Lineage 2.3.4.4b has now been detected on all continents except Australia and Antarctica (8). HPAI incursion risk to Australia has previously been considered low due to the absence of waterfowl species that migrate beyond Australia (9)(Figure 1), as also exemplified from influenza genomic surveillance (10). Still, annually, millions of migratory seabirds and shorebirds migrate from Asia and North America to Australia (Figure 1). Some of these species have been shown to be part of the avian influenza reservoir community (11) and potentially survive and move HPAI viruses (12).

**Figure 1:**
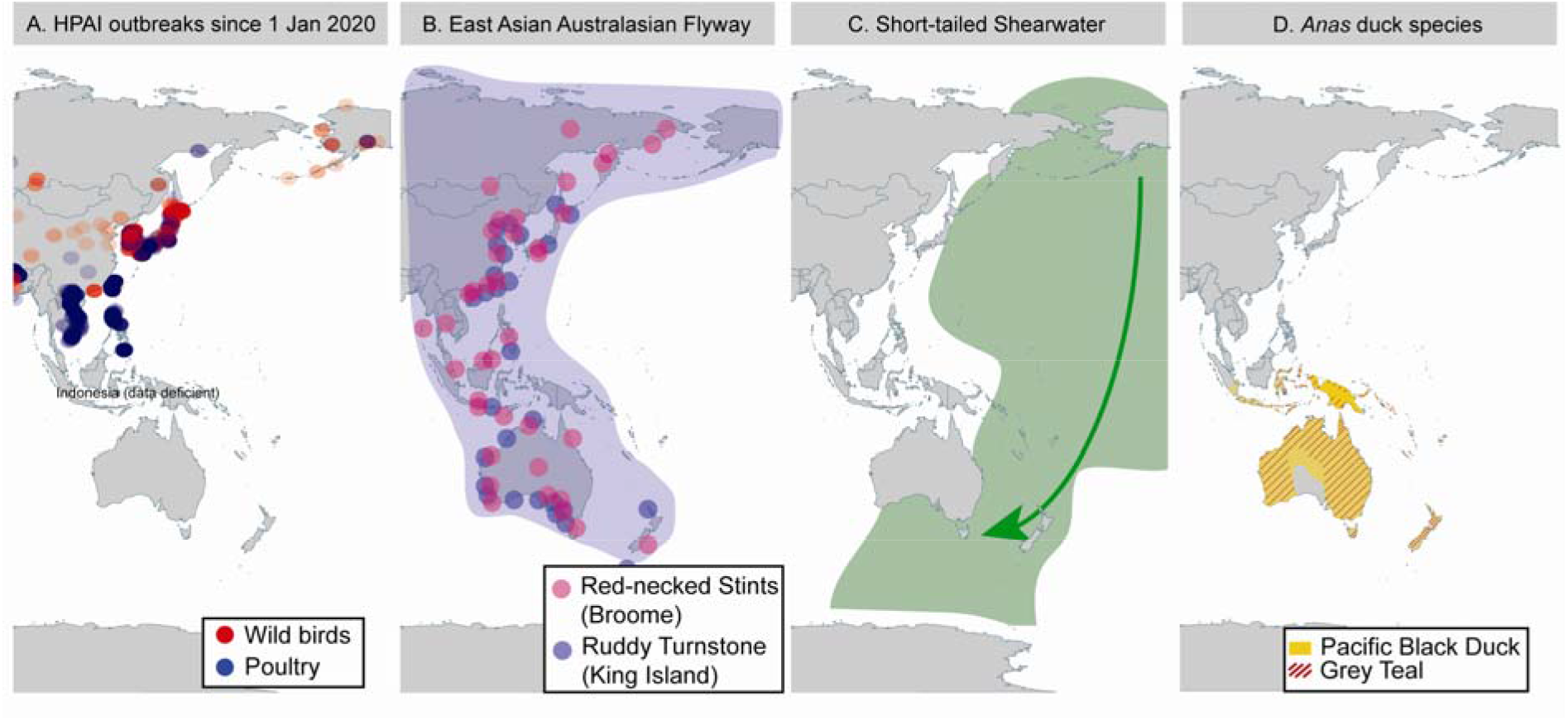
HPAI outbreaks along the East Asian Australasian flyway and distributions of key avian influenza reservoir species in Australia. (A) Outbreaks of HPAI in wild birds (red symbols where intensity reflects number of outbreaks at that location) and poultry (blue symbols) since 1 Jan 2020. Data mined from the World Animal Health Information System of the World Organisation for Animal Health at https://wahis.woah.org/. (B) The East Asian Australasian Flyway utilized by migratory shorebirds. Migratory propensity is exemplified for two populations that we sampled most intensively: Red-necked Stints originally colour-marked in Broome, Western Australia (purple symbols) and Ruddy Turnstones originally marked on King Island, Tasmania (blue symbols). Data extracted from https://www.birdmark.net/. (C) Distribution of Short-tailed Shearwater. Arrow demonstrates southbound migration to Australia occurs from Beringia. (D) Map illustrating the constrating and limited, Australo-papuan distribution of Australian waterfowl using the distribution of Pacific Black Duck (*Anas superciliosa*) and Grey Teal (*Anas gracilis*) as an example. All duck species found in Australia are endemic to Australio-Papuan region and do not migrate to Asia, hence they are likely to play a nominal role in viral incursions from Asia and were therefore not prioritized in this study.

To reveal whether a viral incursion may have occurred in Australia in 2022 with the arrival of wild migratory sea and shorebirds, we investigated 817 migratory birds of the order *Charadriiformes* and *Procelariformes*, in September–December 2022. Specifically, we captured and sampled Short-tailed Shearwaters (n=233) upon their arrival from the northern Pacific to a breeding colony on Philip Island, Victoria, and twelve Asian-breeding migratory shorebird species at major non-breeding sites in Roebuck Bay and 80 mile beach, Western Australia (n=509), and a non-breeding site on King Island, Tasmania (n=75) (Table 1, Figure 1).

**Table 1.**
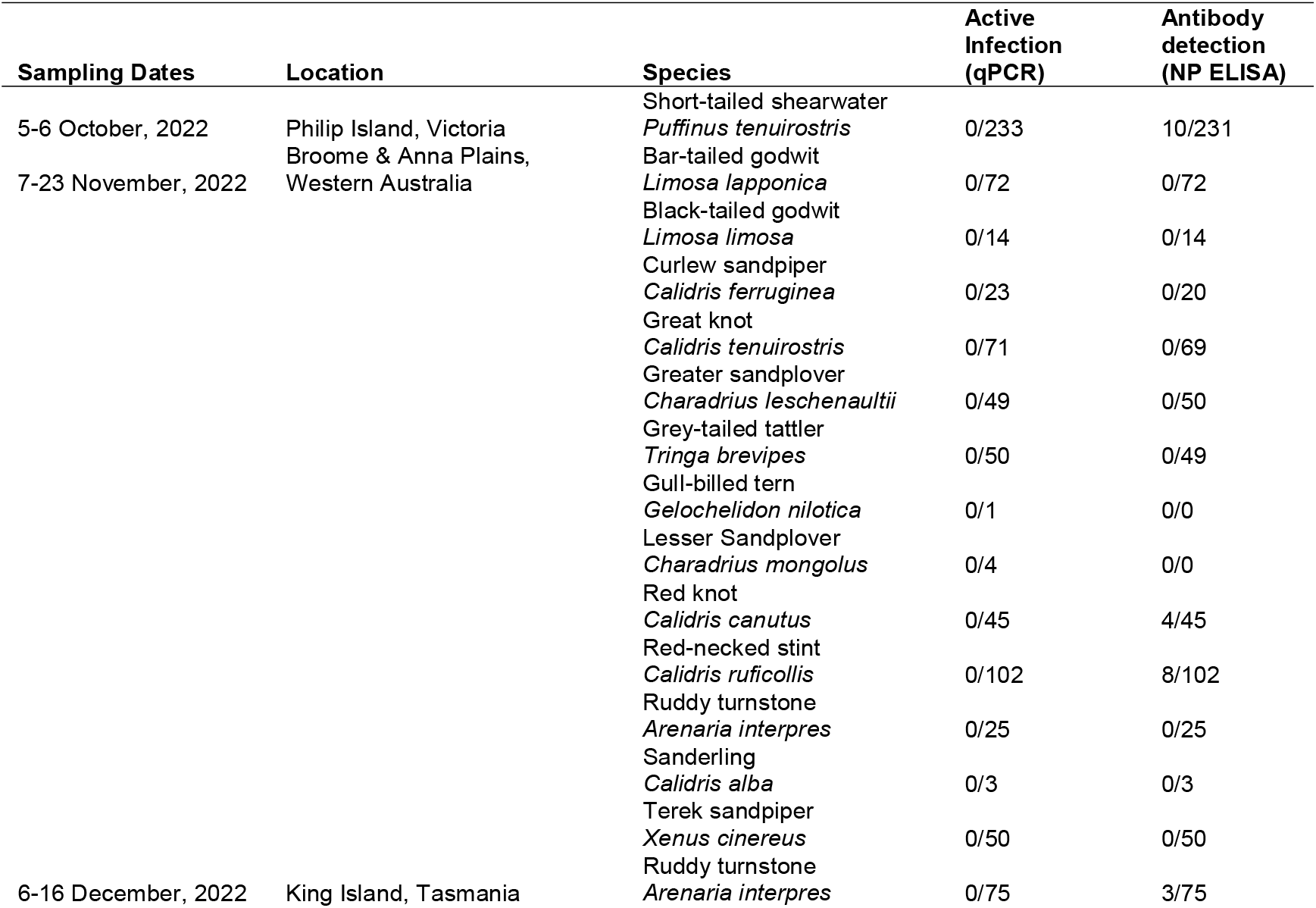
Members of the Charadriiformes and Procellariiformes targeted for active HPAI surveillance following arrival to Australia from migration.

All samples were negative for influenza A virus by qPCR, following (11). Twenty-five serum samples tested positive for anti-NP antibodies using a commercial ELISA (given an S/N cut off of 0.5) (Table 1), which fell within the previously reported seroprevalence of the species that tested positive: Red-necked Stint, Red Knot, Ruddy Turnstone and Short-tailed Shearwater (11). All sera samples positive by anti-NP ELISA were negative on a subsequent hemagglutination inhibition (HI) assay using a lineage 2.3.4.4b candidate vaccine virus A/Astrakhan/3212/2020(H5N8) (13) following (12). A candidate vaccine virus is a 6:2 recombinant virus on an A/Puerto Rico/8/1934(H1N1)(PR8) backbone with the multi-basic cleavage site removed. In addition to the absence of HPAI and antibodies against HPAI lineage 2.3.4.4.b in the sampled migrants, there were neither indications of increased mortality in any wild birds, nor reports of unusual mortality in poultry across Australia.

## Discussion

For Australia as for other regions in the world, HPAI incursion risk hinges on a combination of factors, including wild bird migration, virus pathogenicity in wild birds (notably whether wild birds are able to migrate while infected), and outbreaks and virus circulation in neighbouring regions (particularly at key stopover sites for migratory birds). That there was no incursion of HPAI in Australia in 2022 despite the arrival of millions of migratory birds, the capacity of wild birds to disperse this virus large distances (e.g. (4)), the apparent widening of the virus’ host reservoir beyond waterfowl (7, 8, 14) and and high levels of HPAI activity in Asian countries along the East Asian Australasian flyway (8) is unclear and warrants further investigation.

## Conclusion

Australia will again enter a high-risk period when the major bird migrations into the country take place between August - November 2023. Continued surveillance is critical for early detection and rapid response, and as such we call for enhanced surveillance of Australian wild birds to match heightened incursion risk in the second half of 2023.

## Ethics statement

Capture, banding and sampling was conducted under Victorian Wader Study Group’s ABBBS authority 8001, Deakin University animal ethics committee (B39-2019), Department of Primary Industries and Regional Development WA (20-4-10) and Department of Natural Resources and Environment (5/2019-20).

## Acknowledgements

We wish to acknowledge all those who contributed to bird capture and sample collection including members of Philip Island Nature Parks, Victorian Wader Study Group, Australian Wader Study Group, Broome Bird Observatory. Specifically, we would like to highlight the contributions of Robyn Atkinson, David Boyle, Robert Bush, Tegan Douglas, Richard DuFeu, Teagan Fitzwater, Roz Jessop, Hiske Klaassen, Grace Maglio, Toby Ross, Duncan Sutherland, Teri Visentin and Cassandra Wittwer.

Research in Marcel Klaassen’s lab is undertaken with support from Australian Research Council (ARC) Discovery Project Grant DP19010186. Michelle Wille is funded by an ARC Discovery Early Career Research Award (DE200100977). The WHO Collaborating Centre for Reference and Research on Influenza is funded by the Australian Department of Health.

## Author’s contributions

MW and MK contributed to the conception, data collection, analysis and writing of the article. All authors approve the final manuscript.

